# *Mycobacterium tuberculosis* evasion of Guanylate Binding Protein-mediated host defense in mice requires the ESX1 secretion system

**DOI:** 10.1101/2020.07.27.223362

**Authors:** Andrew J. Olive, Clare M. Smith, Christina E. Baer, Jörn Coers, Christopher M. Sassetti

## Abstract

Cell-intrinsic immune mechanisms control intracellular pathogens that infect eukaryotes. The intracellular pathogen *Mycobacterium tuberculosis* (*Mtb*) evolved to withstand cell-autonomous immunity to cause persistent infections and disease. A potent inducer of cell-autonomous immunity is the lymphocyte-derived cytokine IFNγ. While the production of IFNγ by T cells is essential to protect against *Mtb*, it is not capable of fully eradicating *Mtb* infection. This suggests that *Mtb* evades a subset of IFNγ-mediated antimicrobial responses, yet what mechanisms *Mtb* resists remains unclear. The IFNγ-inducible Guanylate binding proteins (GBPs) are key host defense proteins able to control infections with intracellular pathogens. GBPs were previously shown to directly restrict *Mycobacterium bovis* BCG yet their role during *Mtb* infection has remained unknown. Here, we examine the importance of a cluster of five GBPs on mouse chromosome 3 in controlling Mycobacterial infection. While *M. bovis* BCG is directly restricted by GBPs, we find that the GBPs on chromosome 3 do not contribute to the control of *Mtb* replication or the associated host response to infection. The differential effects of GBPs during *Mtb* versus *M. bovis* BCG infection is at least partially explained by the absence of the ESX1 secretion system from *M. bovis* BCG, since *Mtb* mutants lacking the ESX1 secretion system become similarly susceptible to GBP-mediated immune defense. Therefore, this specific genetic interaction between the murine host and *Mycobacteria* reveals a novel function for the ESX1 virulence system in the evasion of GBP-mediated immunity.

## Introduction

Eukaryotic cells control intracellular pathogens using a variety of cell intrinsic immune pathways (1). These innate mechanisms allow the cell to rapidly detect, target and destroy invading pathogens, preventing the spread of an infection. The immune pathways controlling innate immunity arose early in the evolution of the eukaryota, providing ample time for the selection of pathogens that express mechanisms to bypass cell-autonomous immunity (2). This long-term arms race has produced a myriad of interactions between immune effectors and pathogen countermeasures that determine the outcome of an infection.

*Mycobacterium tuberculosis* (*Mtb*) has become highly adapted to its human host, as it spread globally along with human migrations over tens of thousands of years (3, 4). As much as one-third of the human population has been exposed to *Mtb*, which causes a persistent infection than can last for years, or even decades, despite a robust immune response that eradicates less pathogenic mycobacteria (5). While immunity controls *Mtb* growth and prevents disease in most individuals, a subset will develop active tuberculosis (TB), a disease that kills an estimated 1.8 million each year (6). Disease progression is influenced by a variety of genetic and environment factors, but ultimately is determined by the interplay between host immunity and bacterial virulence systems(5, 7).

In the host, the development of a robust T cell response and the production of the cytokine IFNγ are important for the control TB infection and disease (5, 8). Humans with inherited mutations affecting either the development or expression of this response are highly susceptible to mycobacterial infections including TB (9). This susceptibility is faithfully modeled in mice, where the loss of IFNγ signaling promotes disease by at least two distinct mechanisms (10, 11). IFNγ is an important immunomodulatory cytokine, and its loss results in uncontrolled IL1 production and neutrophil recruitment, driving both bacterial replication and tissue damage (10, 12). Perhaps more importantly, IFNγ stimulates cell-intrinsic immune pathways in phagocytes, which is critical for the control of intracellular bacterial growth (13, 14). Thus, IFNγ is a pleotropic cytokine that controls direct antimicrobial resistance and disease tolerance, both of which are essential to survive *Mtb* infection.

While many of the immunomodulatory effects of IFNγ have been elucidated, the IFNγ-induced factors that control *Mtb* replication remain comparatively obscure. IFNγ-induced oxygen and nitrogen radical generation limits *Mtb* replication in macrophages *ex vivo*, but appear to serve a limited antimicrobial role in the intact animal (14-16). Instead, a subset of IFNγ-inducible cell-intrinsic immune mechanisms, known as Guanylate binding proteins (GBPs), target and disrupt the intracellular niche required for a number of pathogens to grow (2, 17-21). Macrophages lacking GBPs 1, 6,7 or 10 fail to control the growth of *M. bovis* BCG, an attenuated vaccine strain that is closely related to *Mtb (22)*. Similarly, mice lacking GBP1 or the cluster of 5 GBP proteins on chromosome 3 are more susceptible to intravenous BCG challenge (22, 23). While the mechanism of BCG control remains unclear, GBPs are known to bind to pathogen containing vacuoles in other infections and recruit additional effector molecules (17, 24). The outcome of this recognition can be the direct restriction of pathogen growth and alterations in cytokine production. While these observations suggest that GBPs may be an important mediator of IFNγ-mediated control of pathogenic mycobacteria, their role in during *Mtb* infection has remained untested.

BCG was attenuated for use as a vaccine via long term serial passage, and as a result, it interacts with the macrophage quite differently from *Mtb* (25). Most notably, the primary genetic lesion responsible for the attenuation of BCG is a deletion that disrupts the ESX1 type VII secretion system (26). ESX1 is specialized protein secretion complex that contributes to pathogen replication by remodeling its intracellular environment (27). ESX1 is responsible for the disruption of the phagosomal membrane, the activation of multiple cytosolic immune sensing pathways, and stimulation of both cytokine secretion and autophagy (28). How ESX1 alters other aspects of cell-intrinsic bacterial control remains to be determined.

To understand the mechanisms of IFNγ-mediated protection we investigated the role of GBPs during both BCG and *Mtb* infection. Specifically, we examined the function of a cluster of 5 GBP proteins that are encoded in a single locus on mouse chromosome 3 (29). While these GBPs restricted *M. bovis* BCG replication, we found no contribution of these proteins in the control of virulent *Mtb* infection. The discrepant effects of GBPs during BCG and *Mtb* infection could be attributed to differential ESX1 function in these two pathogens. ESX1-deficient strains of *Mtb* were better controlled by GBP-mediated immunity, while GBP deficiency had no effect on either the growth of ESX1 expressing *Mtb* or the immune response to this virulent strain. Together, these observations indicate that the ESX1 system plays an essential role in the evasion of GBP-mediated cell intrinsic immunity.

## Materials and Methods

### Mice

C57BL/6J (Stock # 000664) and IFNγR^-/-^ (Stock # 003288) mice were purchased from the Jackson Laboratory. The Gbp^chr3-/-^ mice were previously described (29). All knockout mice were housed and bred under specific pathogen-free conditions and in accordance with the University of Massachusetts Medical School IACUC guidelines. All animals used for experiments were 6-12 weeks.

### Bacterial Strains

Wild type *M. tuberculosis* strain H37Rv was used for all studies unless indicated. This strain was confirmed to be PDIM-positive. The espA (Rv3616c) deletion strain (Δ*espA*) was a gift from Dr. Sarah Fortune (30). The eccb1 (Rv3869) deletion strain (Δ*eccb1*) was constructed in the H37Rv parental background using the ORBIT method as described previously (31). The live/dead strain was built by transforming the live/dead vector (pmV261 hsp60::mEmerald tetOtetR::TagRFP) into H37Rv and selected with hygromycin. Protein expression was confirmed via fluorescence microscopy and flow cytometry. H37Rv expressing msfYFP has been previously described and the episomal plasmid was maintained with selection in Hygromycin B (50ug/ml) added to the media (32). *Mycobacterium bovis* BCG Danish Strain 1331 (Statens Serum Institute, Copenhagen, Denmark) was used for all BCG infection studies. Prior to infection bacteria were cultured in 7H9 medium containing 10% oleic albumin dextrose catalase growth supplement (OADC) enrichment (Becton Dickinson) and 0.05% Tween 80.

### Mouse Infection

For low dose aerosol infections (50-150 CFU), bacteria were resuspended in phosphate-buffered saline containing tween 80 (PBS-T). Prior to infection bacteria were sonicated then delivered via the respiratory route using an aerosol generation device (Glas-Col). For mixed infections, bacteria were prepared then mixed 1:1 before aerosol infection. To determine CFU, mice were anesthetized via inhalation with isoflurane (Piramal) and euthanized via cervical dislocation, the organs aseptically removed and individually homogenized, and viable bacteria enumerated by plating 10-fold serial dilutions of organ homogenates onto 7H10 agar plates. Plates were incubated at 37C, and bacterial colonies counted after 21 days. Both male and female mice were used throughout the study and no significant differences in phenotypes were observed between sexes.

### Flow Cytometry

Lung tissue was harvested in DMEM containing FBS and placed in C-tubes (Miltenyi). Collagenase type IV/DNaseI was added and tissues were dissociated for 10 seconds on a GentleMACS system (Miltenyi). Tissues were incubated for 30 minutes at 37C with oscillations and then dissociated for an additional 30 seconds on a GentleMACS. Lung homogenates were passaged through a 70-micron filter or saved for subsequent analysis. Cell suspensions were washed in DMEM, passed through a 40-micron filter and aliquoted into 96 well plates for flow cytometry staining. Non-specific antibody binding was first blocked using Fc-Block. Cells were then stained with anti-Ly-6G Pacific Blue, anti-CD11b PE, anti-CD11c APC, anti-Ly-6C APC-Cy7, anti-CD45.2 PercP Cy5.5, anti-CD4 FITC, anti-CD8 APC-Cy7, anti-B220 PE-Cy7 (Biolegend). Live cells were identified using fixable live dead aqua (Life Technologies). For infections with fluorescent H37Rv, lung tissue was prepared as above but no antibodies were used in the FITC channel. All of these experiments contained a non-fluorescent H37Rv infection control to identify infected cells. Cells were stained for 30 minutes at room temperature and fixed in 1% Paraformaldehyde for 60 minutes. All flow cytometry was run on a MACSQuant Analyzer 10 (Miltenyi) and was analyzed using FlowJo_V9 (Tree Star).

### Bone marrow-derived macrophage generation

To generate bone marrow derived macrophages (BMDMs), marrow was isolated from femurs and tibia of age and sex matched mice as previously described (33). Cells were then incubated in DMEM (Sigma) containing 10% fetal bovine serum (FBS) and 20% L929 supernatant. Three days later media was exchanged with fresh media and seven days post-isolation cells were lifted with PBS-EDTA and seeded in DMEM containing 10% FBS for experiments.

### Macrophage Infection

*Mtb* or *Mycobacterium bovis*-BCG were cultured in 7H9 medium containing 10% oleic albumin dextrose catalase growth supplement (OADC) enrichment (Becton Dickinson) and 0.05% Tween 80. Before infection cultures were washed in PBS-T, resuspended in DMEM containing 10%FBS and centrifuged at low speeds to pellet clumps. The supernatant was transferred to a new tube to ensure single cells. Multiplicity of infection (MOI) was determined by optical density (OD) with an OD of 1 being equivalent to 3×10^8^ bacteria per milliliter. Bacteria were added to macrophages for 4 hours then cells were washed with PBS and fresh media was added. At the indicated time points supernatants were harvested for cytokine analysis and the cells were processed for further analysis. For cytokine treatments cells were treated with the indicated concentrations of IFNγ (Peprotech) or vehicle control four hours following infection and maintained in the media throughout the experiment. For CFU experiments at the indicated timepoints 1% saponin was added to each well and lysates were serially diluted in PBS .05% Tween and plated on 7H10 agar and colonies were counted 21-28 days later. For the Live/Dead reporter experiments, Anhydrotetracycline (aTc) (Cayman Chemical) was added to a final concentration of 500ug/ml 24 hours before cells were lifted, fixed in 1% Paraformaldehyde and analyzed on a MacsQuant VYB analyzer.

### Cytokine quantification by ELISA

Murine cytokine concentrations in culture supernatants and cell-free lung homogenates were quantified using commercial enzyme-linked immunosorbent assay (ELISA) kits (R&D). All samples were normalized for total protein content.

### Statistics

GraphPad Prism version 7 was used for all statistical analysis. Unless otherwise indicated one-way ANOVA with a tukey correction was used to compare each condition to each genotype.

## Results

### *Mycobacterium bovis* BCG growth is restricted by the Chromosome 3 Guanylate binding protein cluster

The ten murine GBP genes are encoded in two clusters on chromosomes 3 and 5 (1). To begin to address the potential redundancy between these genes, we used mice lacking the entire chromosome 3 cluster that contains the Gbp1 and 7 genes that were previously implicated in BCG control, along with GBPs 2, 3, and 5 (29). To determine if these GBPs contribute to control of mycobacterial infection, bone marrow-derived macrophages (BMDMs) from wild type and Gbp^chr3-/-^ mice were infected with *Mycobacterium bovis* BCG (Figure 1A). IFNγ-treatment, which induces GBP expression, reduced the intracellular bacterial burden in wild type BMDMs. In contrast, the Gbp^chr3-/-^ deletion significantly reduced the ability of macrophages to control BCG upon IFNγ activation. Thus, GBPs on chromosome 3 contribute to IFNγ-mediated restriction of BCG in macrophages.

**Figure 1.**
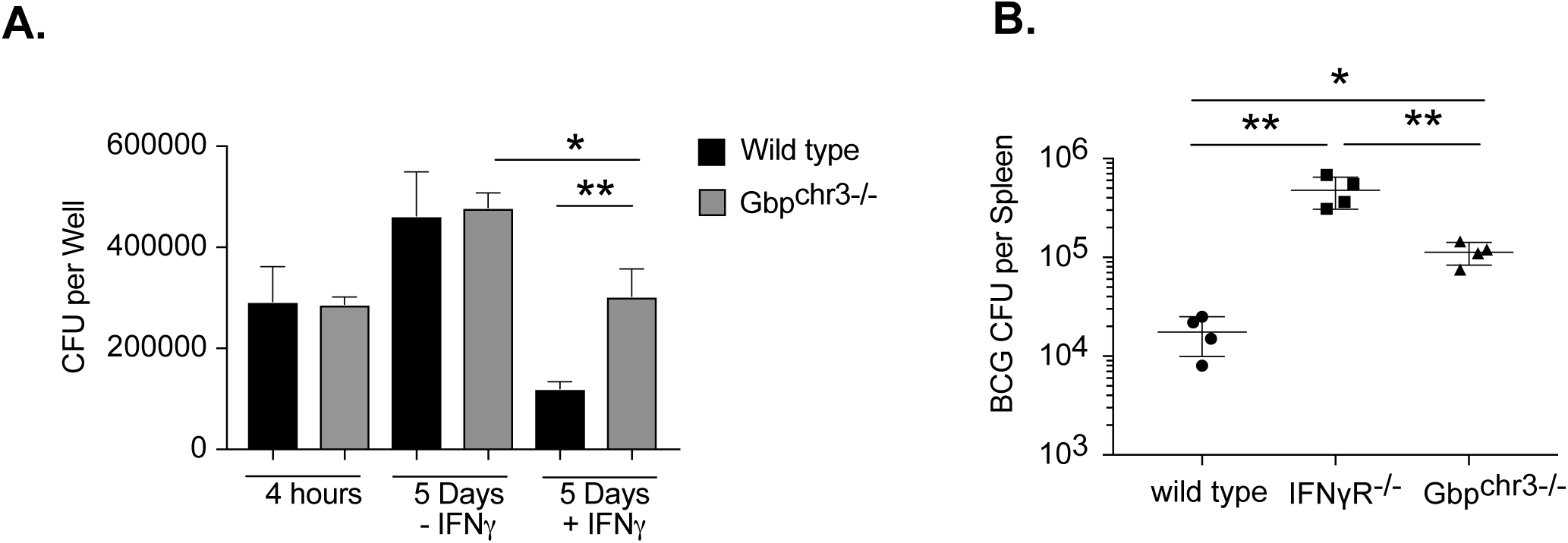
Guanylate binding proteins contribute to control of *Mycobacterium Bovis* BCG infection. **(A)** BMDMs from wild type or Gbp^chr3-/-^ mice were infected with *M. Bovis* BCG for 4 hours then washed with fresh media in the presence or absence of IFNγ. Five days later the macrophages were lysed and serial dilutions were plated to quantify colony forming units (CFU) of viable BCG. Shown is the mean CFU from four biological replicates +/- SD *p<.05 **p<.01. Data are representative of four independent experiments. **(B)** Following IV infection with BCG (1×10^6^ bacteria) the total bacterial burden (expressed in CFU, mean +/- SD) was determined in the spleen of wild type of Gbp^chr3-/-^ mice 50 days following infection. Representative of two independent experiments with 4-5 mice per group, *p<.05 **p<.01.

We next examined if GBPs controlled BCG infection in the context of an intact immune response. Wild type, IFNγR^-/-^ and Gbp^chr3-/-^ mice were infected intravenously with BCG. When CFU were quantified in the spleen 50 days later we observed twenty-fold more BCG in IFNγ mice and five-fold more BCG in Gbp^chr3-/-^ mice compared to wild type controls (Figure 1B). The increased susceptibility of IFNγR^-/-^ compared to Gbp^chr3-/-^ mice indicated that the function of these GBPs accounted for some but not all of the protective effect of IFNγ. Together these data show that the GBPs on chromosome 3 are able to restrict the growth of BCG in IFNγ-stimulated macrophages and in the intact animal, which is consistent with the previously described roles of GBPs in immunity to BCG (22, 23).

### Chromosome 3 GBP cluster does not control *Mycobacterium tuberculosis* growth in macrophages

To determine if the chromosome 3 GBPs restricted the intracellular growth of virulent *Mtb*, in addition to BCG, we quantified the growth of *Mtb* strain H37Rv in BMDMs from wild type, IFNγR^-/-^ and Gbp^chr3-/-^ animals in the presence and absence of IFNγ (Figure 2A). We observed no differences in the uptake between genotypes four hours following infection. IFNγ priming reduced the intracellular *Mtb* growth by 3-4 fold in wild type macrophages, but not IFNγR^-/-^ BMDMs at 5 days post infection (dpi). In contrast to BCG, we observed no difference in *Mtb* growth in Gbp^chr3-/-^ BMDMs compared to wild type macrophages in the presence or absence of IFNγ.

**Figure 2.**
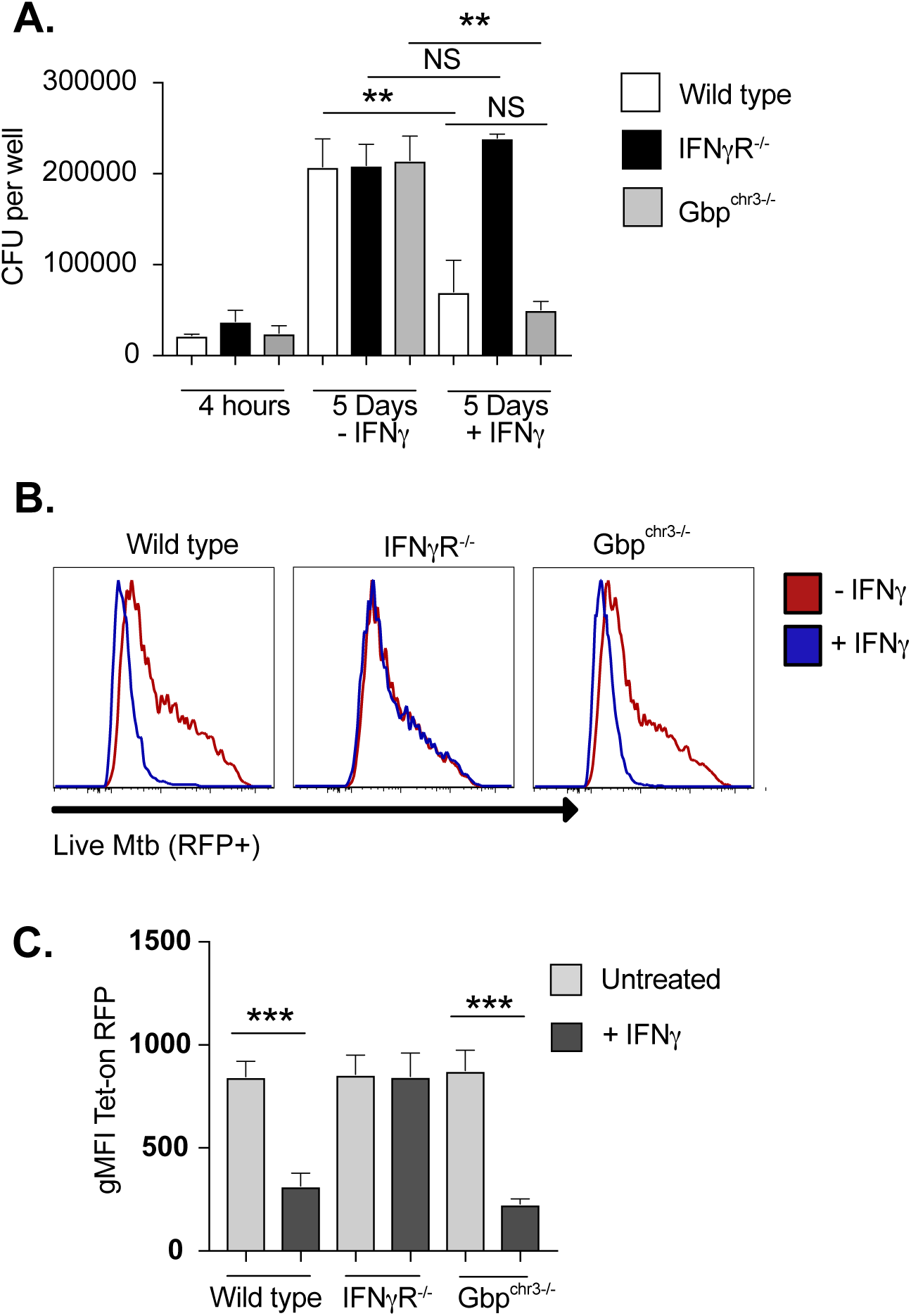
*Mycobacterium tuberculosis* is resistant to chromosome 3 GBP-mediated control in macrophages. **(A)** BMDMs from wild type, IFNγR^/-^ or Gbp^chr3-/-^ mice were infected with *M. tuberculosis* for 4 hours then washed with fresh media in the presence or absence of IFNγ. Five days later the macrophages were lysed, serial dilutions were plated to quantify colony forming units (CFU) of viable *M. tuberculosis*. Shown is the mean CFU from four biological replicates +/- SD *p<.05 **p<.01. Data are representative of five independent experiments. **(B)** and **(C)** BMDMs from wild type, IFNγR^/-^ or Gbp^chr3-/-^ mice were infected with the live/dead reporter *M. tuberculosis* for 4 hours then washed with fresh media in the presence or absence of IFNγ. Four days later aTc was added to each well at a final concentration of 500ng/ml. The following day infected cells were lifted and the fluorescence intensity of the inducible Tet-on TagRFP was determined by flow cytometry. Cells were gated on live and infected (mEmerald+) cells. **(B)** A representative histogram of tagRFP fluorescence is shown. **(C)** Shown is the mean fluorescence intensity of tagRFP for three biological replicates. These data are representative of three independent experiments. ***p<.001.

To ensure that the relatively insensitive CFU-based intracellular growth assay did not mask subtle effects of GBP expression on bacterial fitness, we used a fluorescent live/dead reporter as an orthologous method to measure *Mtb* intracellular viability by flow cytometry. This reporter expresses a constitutive mEmerald and an anhydrotetracycline (aTc)-inducible tagRFP. BMDMs from wild type, IFNγR^-/-^ and Gbp^chr3-/-^ mice were infected with the Live/Dead *Mtb* reporter and left untreated or were stimulated with IFNγ. Four days later aTc was added to induce tagRFP expression in viable intracellular bacteria. The following day, cells were analyzed by flow cytometry and the mean fluorescence intensity (MFI) of tagRFP in infected mEmerald+ cells was quantified (Figure 2B and 2C). Similar to the CFU analysis, IFNγ activation reduced the intensity of tagRFP to a similar extent in both wild type and Gbp^chr3-/-^ BMDMs, and this reduction depended on the IFNγ receptor. Thus, we were not able to detect a role for the chromosome 3 GBP cluster in the IFNγ mediated restriction of virulent *Mtb* in macrophages.

### Chromosome 3 GBPs have no effect on *Mtb* infection in intact mice

To assess the role of the chromosome 3 GBPs in the more complex setting of the intact animal, we infected wild type, IFNγR^-/-^ and Gbp^chr3-/-^ mice with *Mtb* by low dose aerosol and quantified bacteria in the lungs at four and five weeks following infection (Figure 3A and 3B). IFNγR^-/-^ mice harbored more *Mtb* than wild type controls at both time points in the lung and the spleen. In contrast, the *Mtb* burdens in Gbp^chr3-/-^ mice were indistinguishable from wild type animals at all timepoints. At 110 days after infection (Figure 3A and 3B), we continued to find no difference in CFU in the lungs between wild type and GBP^chr3-/-^ mice. IFNγR^-/-^ mice required euthanasia within six weeks of infection and were not included in this late time point.

**Figure 3.**
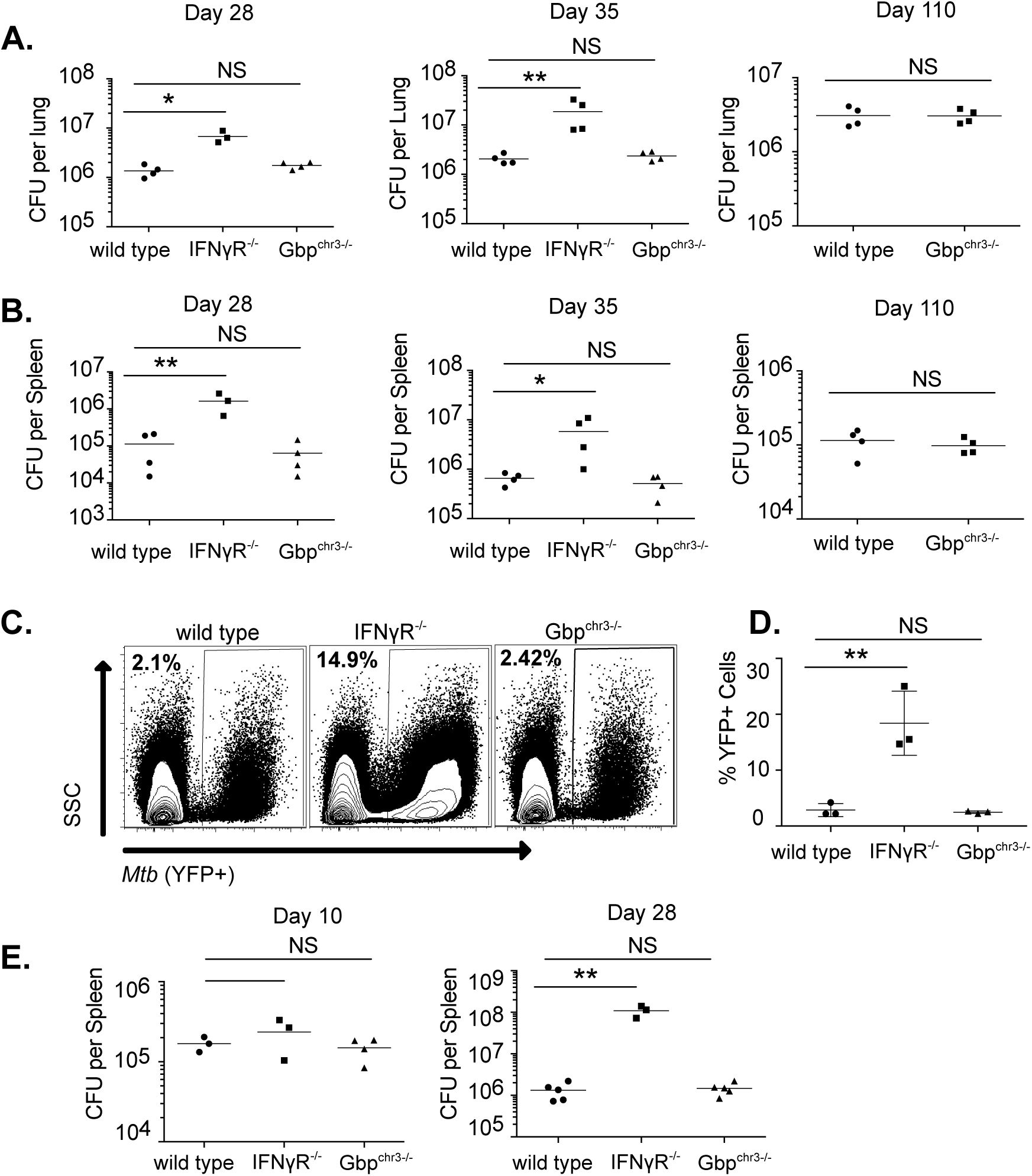
Mice lacking chromosome 3 GBPs control *M. tuberculosis* infection. Following low-dose aerosol infection with H37Rv (day 0 of ∼50-150 CFU), total bacteria (expressed in CFU, mean +/- SD) was determined in the lungs **(A)** or the spleen **(B)** of wild type, IFNγR^-/-^ or Gbp^chr3-/-^ mice at the indicated time points with four mice per group. 28 days following low-dose aerosol infection with sfYFP H37Rv, infected cells in the lungs of wild type, IFNγR^-/-^ or Gbp^chr3-/-^ mice were quantified by flow cytometry. **(C)** Shown is a representative flow cytometry plot of the infected cells (YFP positive) that are gated on live, single cells and **(D)** the mean percent of YFP positive cells in the lungs of infected animals. **(E)** Following intravenous infection with H37Rv (1×10^6^ per mouse), total bacteria were determined in the spleen of wild type, IFNγR^-/-^ or Gbp^chr3-/-^ mice at the indicated time points with 3-5 mice per group. All data are representative of 3 independent experiments.

To examine more subtle differences in the extent of infection, wild type, IFNγR^-/-^ and Gbp^chr3-/-^ mice were infected with a well-characterized fluorescent H37Rv strain by low-dose aerosol (32). Twenty-eight days later the number of YFP+ cells in the lung environment were determined by flow cytometry (Figure 3C). Similar to our CFU results, we found IFNγR mice contained over 10 times more infected cells than wild type mice while Gbp^chr3-/-^ mice showed no significant difference in the number of infected cells per lung (Figure 3D).

We next examined if there were differences in Gbp^chr3-/-^ mice infected intravenously since this route matched the BCG infections where the role of GBPs was evident (Figure 1B and (22)). Following intravenous infection with 10^6^ *Mtb*, bacterial levels in the spleen were quantified 10 and 28 days later. Similar to the aerosol infection results, we saw no difference between Gbp^chr3-/-^ and wild type animals, while IFNγR^-/-^ animals had 10 times more *Mtb* growth in the spleen (Figure 3E). Thus, unlike the loss of IFNγR, the loss of chromosome 3 GBPs does not affect *Mtb* growth in the lungs or spleen, regardless of infection route, suggesting no major function of these five GBPs in controlling antimicrobial resistance to *Mtb* in mice.

### Gbp^chr3-/-^ mice continue to regulate inflammatory responses to *Mtb* infection

IFNγ protects against disease both by restricting bacterial growth and by inhibiting tissue-damaging inflammation (11). To determine if the chromosome 3 GBPs contribute to the latter immunoregulatory function of IFNγ, we profiled the immune responses of infected macrophages and mice. Wild type and GBP^chrm3-/-^ BMDMs were treated with IFNγ then infected with *Mtb* and the following day supernatants were harvested. IFNγ signaling showed the expected inhibitory effect on IL-1β secretion, but the chromosome3 Gbp deletion had no effect on either cytokine (12). We next profiled the host response in the lungs of mice to determine if chromosome 3 GBPs modulated the recruitment of immune cells or the production of cytokines during *Mtb* infection. Wild type, IFNγR^-/-^ and Gbp^chr3-/-^ mice were infected with *Mtb* and four weeks later we quantified the immune populations in the lungs by flow cytometry (Figure 4B). While we observed the previously described increase in neutrophil recruitment in IFNγR^-/-^ mice, we observed no differences in any myeloid or lymphoid derived cells that were examined between Gbp^chr3-/-^ and wild type mice (10). We also quantified IL1β and TNFα in the lungs from these infected animals (Figure 4C). The production of these cytokines in wild type and GBP^chrm3-/-^ mice were indistinguishable while IFNγR^-/-^ mice showed an increase in IL1β. Thus, we were unable to detect a role for GBPs on chromosome 3 in controlling the host response during virulent *Mtb* infection.

**Figure 4.**
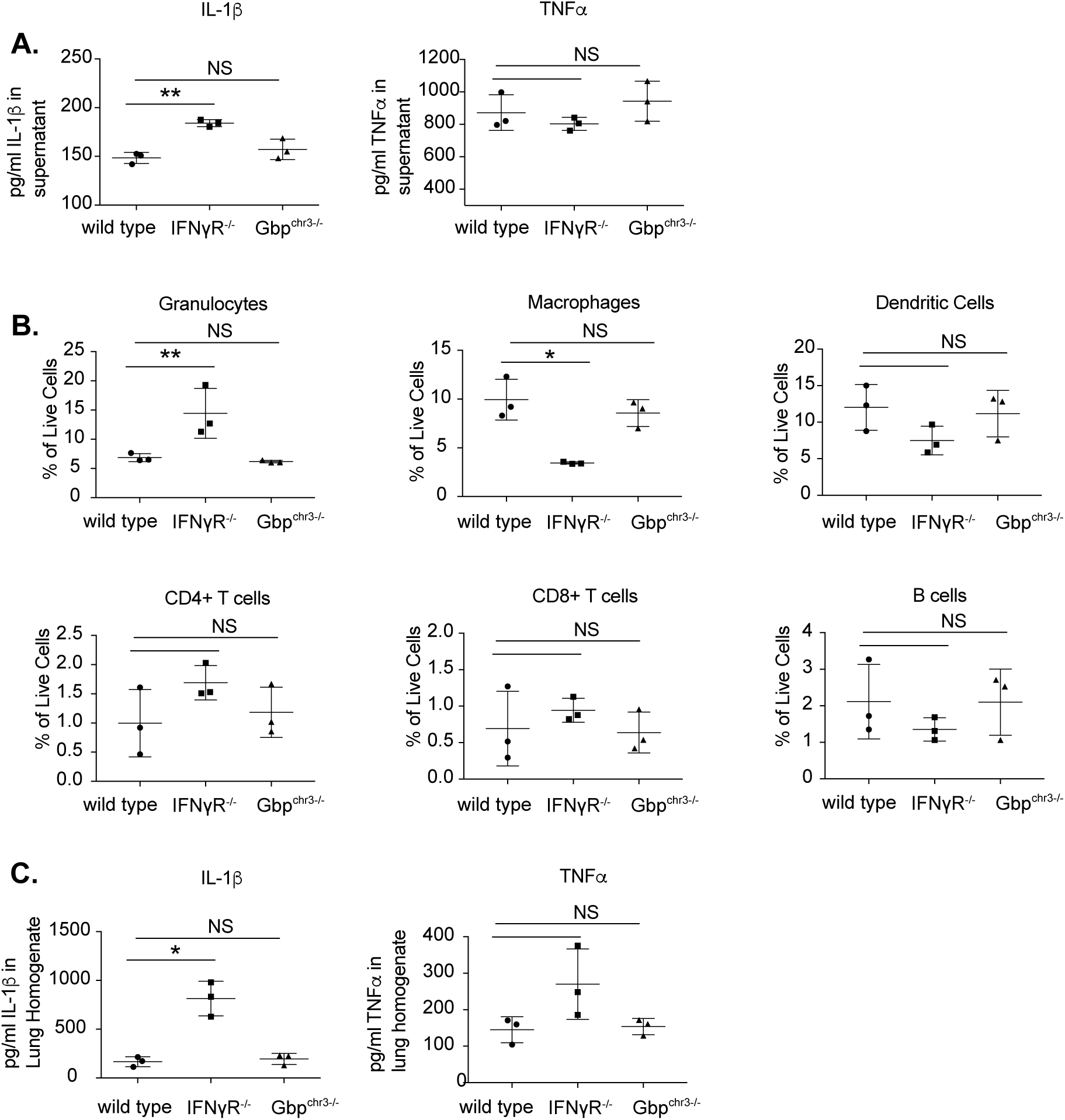
Chromosome 3 GBPs do not control the host response to *M. tuberculosis* infection. **(A)** BMDMs from wild type, IFNγR^/-^ or Gbp^chr3-/-^ mice were stimulated with IFNγ overnight then infected with *Mtb* for 4 hours then washed with fresh media. 18 hours later, supernatants were harvested and the levels of IL1β and TNFα were quantified in the supernatants by ELISA. Shown is the mean of four biological replicates normalized to a standard curve +/- SD *p<.05. **(B)** and **(C)** 28 days following low-dose aerosol infection with H37Rv (day 0 of ∼50-100 CFU), the lungs of wild type, IFNγR^/-^, and Gbp^chr3-/-^ mice were harvested and the immune cell populations were quantified by flow cytometry and cytokines were quantified by ELISA. For **(B)**, we determined the % of live cells for Granulocytes (CD45^+^ CD11b^+^ Ly6G^+^), Macrophages (CD45^+^ CD11b^+^ Ly6G^-^), Dendritic Cells (CD45^+^ CD11b^-^ Ly6G^-^ CD11c^+^), CD4^+^ T cells (CD45^+^ CD4^+^), CD8^+^ T cells (CD45^+^ CD8^+^) and B cells (CD45^+^ B220^+^). For **(C)** Cytokines quantified from lung homogenates. Shown is the mean of three mice per group +/- SD *p<.05. These data are representative of 3-4 independent experiments with similar results.

### ESX1 is required for *Mtb* to evade GBPs of chromosome 3

The reduced virulence of BCG compared to *Mtb* has been largely attributed to the loss of ESX1 function in BCG. As a result, we hypothesized ESX1 function in virulent *Mtb* provides resistance to GBP-mediated immunity, and its loss renders BCG susceptible to this mechanism. To test this hypothesis, we determined whether abrogation of ESX1 function in *Mtb* would result in a strain that was susceptible to GBP-mediated control. Wild type and Gbp^chr3-/-^ macrophages were infected with H37Rv or two isogenic mutants each lacking a gene that is necessary for ESX1 function, Δ*espA* or Δ*eccb1*. We observed that ESX1 mutants displayed reduced intracellular growth compared to H37Rv in both IFNγ stimulated and unstimulated wild type macrophages, confirming the attenuation of these mutants (Figure 5A and 5B). When we compared wild type and Gbp^chr3-/-^ BMDMs, we found that the loss of GBP function had no effect on H37Rv, while it significantly reduced the ability of IFNγ to control the growth of both ESX1 mutants.

**Figure 5.**
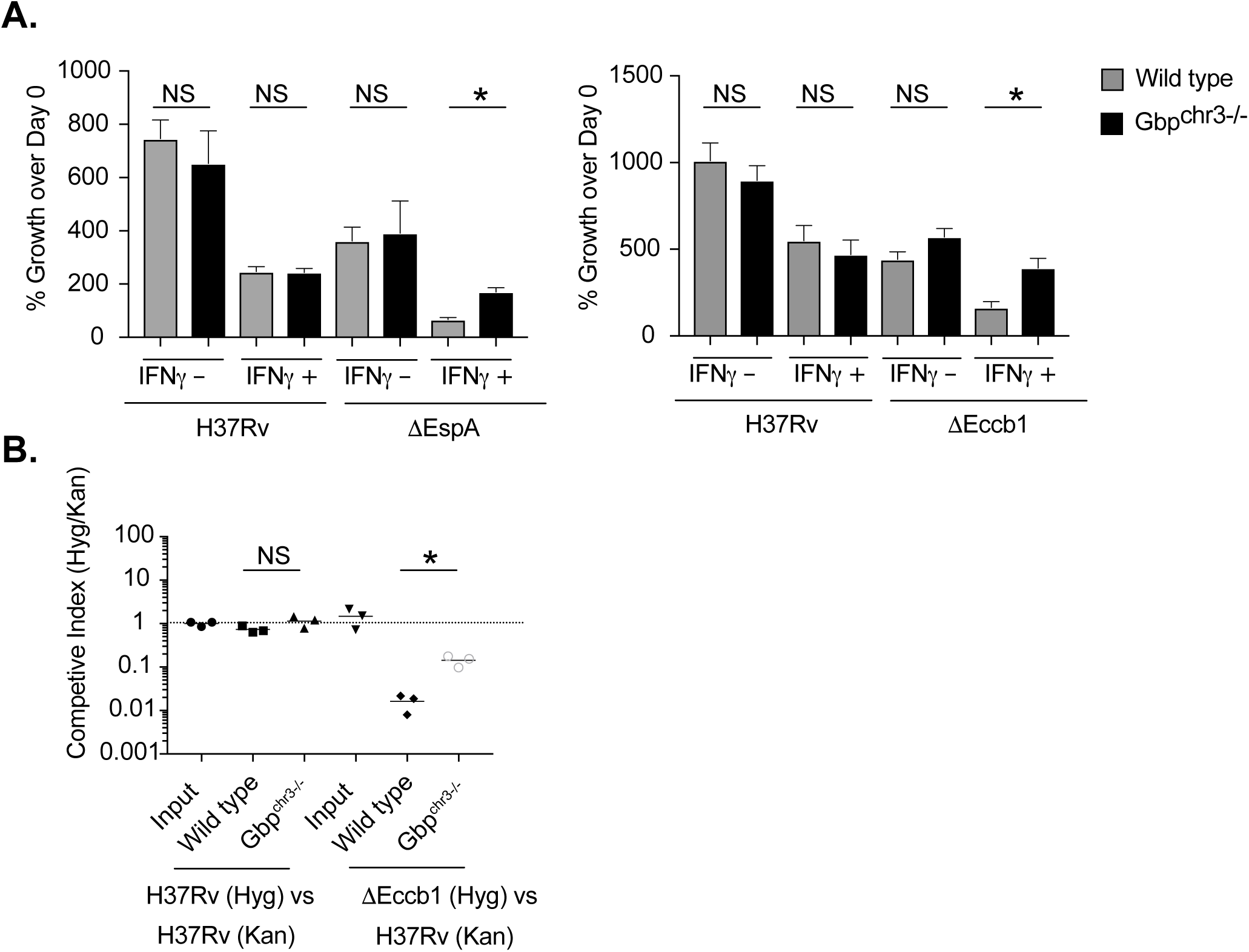
*M. tuberculosis* ESX1 mutants are susceptible to GBP-mediated control. **(A)** BMDMs from wild type or Gbp^chr3-/-^mice were infected with *M. tuberculosis* H37Rv, ΔespA (Left) or Δeccb1 (Right) for 4 hours then washed with fresh media in the presence or absence of IFNγ. Five days later the macrophages were lysed and serial dilutions were plated to quantify colony forming units (CFU) of viable *M. tuberculosis*. Shown is the mean CFU from four biological replicates +/- SD *p<.05 **p<.01 by two-tailed t-test. Data are representative of two independent experiments with similar results. (B) Following low-dose aerosol infection with a 1:1 mixed infection of either H37Rv (Hyg):H37Rv (Kan) or Δeccb1 (Hyg): H37Rv (Kan) (day 0 of ∼100-200 CFU), total bacteria in the lungs that were either Kan or Hyg resistant were quantified. The competitive index was calculated (Hyg CFU/Kan CFU) and is shown as the mean for 3 independent mice for each genotype combination. The data are representative of two independent experiments with similar results.

To confirm the specific effect of chromosome 3 GBPs on ESX1 mutants in the setting of intact immunity, we conducted a competitive infection of wild type and Gbp^chr3-/-^ mice with an equivalent number of H37Rv and Δ*eccb1* bacteria. Four weeks later we quantified the competitive index of each bacterial strain in the lungs (Figure 5C). As anticipated, the ratio of two differentially marked H37Rv strains stayed constant throughout the infection. While selection in wild type mice resulted in almost 100-fold underrepresentation of the Δ*eccb1* mutant compared H37Rv, the fitness defect of the Δ*eccB1* mutant was significantly reduced in the GBP^chr3-/-^ mice (Figure 5C). These findings demonstrate that ESX1 deficiency is sufficient to render *Mtb* susceptible to GBP-mediated immunity.

## Discussion

Understanding the immune mechanisms that restrict the intracellular growth of *Mtb* is essential for the rational design of interventions. Initial observations demonstrating a role for GBPs in the control of *BCG* growth suggested that this pathway might represent an important component of IFNγ-mediated immunity to *Mtb* (22, 23). However, while we were able detect the previously-described role for the chromosome 3 GBPs in immunity to BCG, we found that these proteins have no effect during *Mtb* infection. By attributing this difference to the ESX1 locus that is present in *Mtb* but not BCG, we discovered a specific role for ESX1 in overcoming GBP-mediated defenses.

Our findings question the importance of GBPs in the control of ESX1-expressing *Mtb*. While our results are strictly in the mouse model, evidence in humans also suggests GBPs may not effectively control TB progression. For example, the high expression of a subset of GBPs is predictive of patients that are more likely to progress to active disease (34). However, it is important to note that it remains possible that the chromosome 5 GBPs, or human-specific GBP functions, can overcome ESX1-mediated bacterial defenses. In addition, our findings do not rule out an important role for GBPs in resistance to non-tuberculous mycobacteria (NTM). While all mycobacteria express ESX paralogs, many pathogenic NTM do not possess a clear ortholog of ESX1, suggesting that they may remain susceptible to GBP immunity (35).

How the ESX1 type VII secretion system allows *Mtb* to evade restriction by GBPs remains to be investigated. To date, the type III secretion system effector protein IpaH9.8 from *Shigella flexneri* is the only other described GBP antagonist (36-39). In the cytosol of mammalian cells, where *S. flexneri* replicates, IpaH9.8 targets GBP1 for degradation thereby interfering with direct GBP binding to the *Shigella* outer membrane, a process by which GBP1 disrupts the function of a membrane-bound *Shigella* virulence factors required for actin-based motility and bacterial dissemination (36-38, 40). Similar to IpaH9.8, ESX1 and its substrates may act as direct GBP antagonist to prevent binding to *Mycobacteria*-containing vacuoles. Alternatively, ESX1 may control the evasion of antimicrobial mechanisms that occur subsequent to GBP translocation to *Mycobacteria*-containing vacuoles. Future work will need to dissect these possible mechanisms for ESX1-mediated antagonism of GBP function.

While IFNγ is unquestionably important to survive *Mtb* infection, the IFNγ-mediated pathways that directly control the intracellular replication of *Mtb* remain surprisingly unclear. Our results add to a growing list of direct antimicrobial pathways that are ineffective during *Mtb* infection. The IFNγ-mediated production of nitric oxide, reactive oxygen species and itaconate kills many pathogens; however, these mechanisms appear to play a small role in directly controlling *Mtb* growth *in vivo* (32, 33, 41). Instead, these mediators are required to inhibit persistent inflammation and to prevent disease progression. In addition, the IFNγ-regulated immunity related GTPases (IRG) family protein, Irgm1, was originally described to target the *Mtb* containing vacuole to control pathogen growth, but recent evidence has questioned whether Irgm1 targets *Mtb* phagosomes (42, 43). Instead, the lymphopenia observed in Irgm1-deficient mice may be predominantly responsible for their susceptibility to *Mtb* similar to other pathogens like *Chlamydia trachomatis* (44, 45). While other pathways have been suggested to play a role in IFNγ-mediated control, including the production of Cathepsins, the role of these mediators in protection *in vivo* remains unclear (46). Overall, our findings add GBP-mediated immunity to the list of IFNγ dependent host defense programs to which *Mtb* has evolved specific counter immune mechanisms blunting the effectiveness of these antimicrobial effectors and thus driving pathogen persistence and disease.

## Acknowledgements

We thank Dr. Masahiro Yamamoto for generously sharing Gbp^chr3-/-^ mice for these studies, and Christina Baer for the development of fluorescent Mtb reporters. This work was supported by grants from the NIH to CMS (AI132130) and the Arnold and Mabel Beckman Foundation to AJO.

## References

1. Randow F, MacMicking JD, James LC. 2013. Cellular self-defense: how cell-autonomous immunity protects against pathogens. Science 340:701–6.

2. Tretina K, Park ES, Maminska A, MacMicking JD. 2019. Interferon-induced guanylate-binding proteins: Guardians of host defense in health and disease. J Exp Med 216:482–500.

3. Saelens JW, Viswanathan G, Tobin DM. 2019. Mycobacterial Evolution Intersects With Host Tolerance. Front Immunol 10:528.

4. Brites D, Gagneux S. 2015. Co-evolution of Mycobacterium tuberculosis and Homo sapiens. Immunol Rev 264:6–24.

5. Nunes-Alves C, Booty MG, Carpenter SM, Jayaraman P, Rothchild AC, Behar SM. 2014. In search of a new paradigm for protective immunity to TB. Nat Rev Microbiol 12:289–99.

6. Zumla A, George A, Sharma V, Herbert RH, Baroness Masham of I, Oxley A, Oliver M. 2015. The WHO 2014 global tuberculosis report--further to go. Lancet Glob Health 3:e10–2.

7. Olive AJ, Sassetti CM. 2016. Metabolic crosstalk between host and pathogen: sensing, adapting and competing. Nat Rev Microbiol 14:221–34.

8. Ernst JD. 2012. The immunological life cycle of tuberculosis. Nat Rev Immunol 12:581–91.

9. Bustamante J, Boisson-Dupuis S, Abel L, Casanova JL. 2014. Mendelian susceptibility to mycobacterial disease: genetic, immunological, and clinical features of inborn errors of IFN-gamma immunity. Semin Immunol 26:454–70.

10. Nandi B, Behar SM. 2011. Regulation of neutrophils by interferon-gamma limits lung inflammation during tuberculosis infection. J Exp Med 208:2251–62.

11. Olive AJ, Sassetti CM. 2018. Tolerating the Unwelcome Guest; How the Host Withstands Persistent Mycobacterium tuberculosis. Front Immunol 9:2094.

12. Mishra BB, Rathinam VA, Martens GW, Martinot AJ, Kornfeld H, Fitzgerald KA, Sassetti CM. 2013. Nitric oxide controls the immunopathology of tuberculosis by inhibiting NLRP3 inflammasome-dependent processing of IL-1beta. Nat Immunol 14:52–60.

13. Braverman J, Stanley SA. 2017. Nitric Oxide Modulates Macrophage Responses to Mycobacterium tuberculosis Infection through Activation of HIF-1alpha and Repression of NF-kappaB. J Immunol 199:1805–1816.

14. Denis M. 1991. Interferon-gamma-treated murine macrophages inhibit growth of tubercle bacilli via the generation of reactive nitrogen intermediates. Cell Immunol 132:150–7.

15. Darwin KH, Ehrt S, Gutierrez-Ramos JC, Weich N, Nathan CF. 2003. The proteasome of Mycobacterium tuberculosis is required for resistance to nitric oxide. Science 302:1963–6.

16. Nambi S, Long JE, Mishra BB, Baker R, Murphy KC, Olive AJ, Nguyen HP, Shaffer SA, Sassetti CM. 2015. The Oxidative Stress Network of Mycobacterium tuberculosis Reveals Coordination between Radical Detoxification Systems. Cell Host Microbe 17:829–37.

17. Pilla-Moffett D, Barber MF, Taylor GA, Coers J. 2016. Interferon-Inducible GTPases in Host Resistance, Inflammation and Disease. J Mol Biol 428:3495–513.

18. Gomes MTR, Cerqueira DM, Guimaraes ES, Campos PC, Oliveira SC. 2019. Guanylate-binding proteins at the crossroad of noncanonical inflammasome activation during bacterial infections. J Leukoc Biol 106:553–562.

19. Man SM, Place DE, Kuriakose T, Kanneganti TD. 2017. Interferon-inducible guanylate-binding proteins at the interface of cell-autonomous immunity and inflammasome activation. J Leukoc Biol 101:143–150.

20. Praefcke GJK. 2018. Regulation of innate immune functions by guanylate-binding proteins. Int J Med Microbiol 308:237–245.

21. Santos JC, Broz P. 2018. Sensing of invading pathogens by GBPs: At the crossroads between cell-autonomous and innate immunity. J Leukoc Biol 104:729–735.

22. Kim BH, Shenoy AR, Kumar P, Das R, Tiwari S, MacMicking JD. 2011. A family of IFN-gamma-inducible 65-kD GTPases protects against bacterial infection. Science 332:717–21.

23. Marinho FV, Fahel JS, de Araujo A, Diniz LTS, Gomes MTR, Resende DP, Junqueira-Kipnis AP, Oliveira SC. 2020. Guanylate binding proteins contained in the murine chromosome 3 are important to control mycobacterial infection. J Leukoc Biol doi: 10.1002/JLB.4MA0620-526RR.

24. Coers J. 2017. Sweet host revenge: Galectins and GBPs join forces at broken membranes. Cell Microbiol 19.

25. Moliva JI, Turner J, Torrelles JB. 2017. Immune Responses to Bacillus Calmette-Guerin Vaccination: Why Do They Fail to Protect against Mycobacterium tuberculosis? Front Immunol 8:407.

26. Tiwari S, Casey R, Goulding CW, Hingley-Wilson S, Jacobs WR, Jr. 2019. Infect and Inject: How Mycobacterium tuberculosis Exploits Its Major Virulence-Associated Type VII Secretion System, ESX-1. Microbiol Spectr 7.

27. Stanley SA, Raghavan S, Hwang WW, Cox JS. 2003. Acute infection and macrophage subversion by Mycobacterium tuberculosis require a specialized secretion system. Proc Natl Acad Sci U S A 100:13001–6.

28. Watson RO, Manzanillo PS, Cox JS. 2012. Extracellular M. tuberculosis DNA targets bacteria for autophagy by activating the host DNA-sensing pathway. Cell 150:803–15.

29. Yamamoto M, Okuyama M, Ma JS, Kimura T, Kamiyama N, Saiga H, Ohshima J, Sasai M, Kayama H, Okamoto T, Huang DC, Soldati-Favre D, Horie K, Takeda J, Takeda K. 2012. A cluster of interferon-gamma-inducible p65 GTPases plays a critical role in host defense against Toxoplasma gondii. Immunity 37:302–13.

30. Garces A, Atmakuri K, Chase MR, Woodworth JS, Krastins B, Rothchild AC, Ramsdell TL, Lopez MF, Behar SM, Sarracino DA, Fortune SM. 2010. EspA acts as a critical mediator of ESX1-dependent virulence in Mycobacterium tuberculosis by affecting bacterial cell wall integrity. PLoS Pathog 6:e1000957.

31. Murphy KC, Nelson SJ, Nambi S, Papavinasasundaram K, Baer CE, Sassetti CM. 2018. ORBIT: a New Paradigm for Genetic Engineering of Mycobacterial Chromosomes. mBio 9.

32. Mishra BB, Lovewell RR, Olive AJ, Zhang G, Wang W, Eugenin E, Smith CM, Phuah JY, Long JE, Dubuke ML, Palace SG, Goguen JD, Baker RE, Nambi S, Mishra R, Booty MG, Baer CE, Shaffer SA, Dartois V, McCormick BA, Chen X, Sassetti CM. 2017. Nitric oxide prevents a pathogen-permissive granulocytic inflammation during tuberculosis. Nat Microbiol 2:17072.

33. Olive AJ, Smith CM, Kiritsy MC, Sassetti CM. 2018. The Phagocyte Oxidase Controls Tolerance to Mycobacterium tuberculosis Infection. J Immunol doi: 10.4049/jimmunol.1800202.

34. Zak DE, Penn-Nicholson A, Scriba TJ, Thompson E, Suliman S, Amon LM, Mahomed H, Erasmus M, Whatney W, Hussey GD, Abrahams D, Kafaar F, Hawkridge T, Verver S, Hughes EJ, Ota M, Sutherland J, Howe R, Dockrell HM, Boom WH, Thiel B, Ottenhoff THM, Mayanja-Kizza H, Crampin AC, Downing K, Hatherill M, Valvo J, Shankar S, Parida SK, Kaufmann SHE, Walzl G, Aderem A, Hanekom WA, Acs, groups GCcs. 2016. A blood RNA signature for tuberculosis disease risk: a prospective cohort study. Lancet 387:2312–2322.

35. Johansen MD, Herrmann JL, Kremer L. 2020. Non-tuberculous mycobacteria and the rise of Mycobacterium abscessus. Nat Rev Microbiol 18:392–407.

36. Wandel MP, Pathe C, Werner EI, Ellison CJ, Boyle KB, von der Malsburg A, Rohde J, Randow F. 2017. GBPs Inhibit Motility of Shigella flexneri but Are Targeted for Degradation by the Bacterial Ubiquitin Ligase IpaH9.8. Cell Host Microbe 22:507–518 e5.

37. Piro AS, Hernandez D, Luoma S, Feeley EM, Finethy R, Yirga A, Frickel EM, Lesser CF, Coers J. 2017. Detection of Cytosolic Shigella flexneri via a C-Terminal Triple-Arginine Motif of GBP1 Inhibits Actin-Based Motility. mBio 8.

38. Li P, Jiang W, Yu Q, Liu W, Zhou P, Li J, Xu J, Xu B, Wang F, Shao F. 2017. Ubiquitination and degradation of GBPs by a Shigella effector to suppress host defence. Nature 551:378–383.

39. Ji C, Du S, Li P, Zhu Q, Yang X, Long C, Yu J, Shao F, Xiao J. 2019. Structural mechanism for guanylate-binding proteins (GBPs) targeting by the Shigella E3 ligase IpaH9.8. PLoS Pathog 15:e1007876.

40. Kutsch M, Sistemich L, Lesser CF, Goldberg MB, Herrmann C, Coers J. 2020. Direct binding of polymeric GBP1 to LPS disrupts bacterial cell envelope functions. EMBO J 39:e104926.

41. Nair S, Huynh JP, Lampropoulou V, Loginicheva E, Esaulova E, Gounder AP, Boon ACM, Schwarzkopf EA, Bradstreet TR, Edelson BT, Artyomov MN, Stallings CL, Diamond MS. 2018. Irg1 expression in myeloid cells prevents immunopathology during M. tuberculosis infection. J Exp Med 215:1035–1045.

42. MacMicking JD, Taylor GA, McKinney JD. 2003. Immune control of tuberculosis by IFN-gamma-inducible LRG-47. Science 302:654–9.

43. Springer HM, Schramm M, Taylor GA, Howard JC. 2013. Irgm1 (LRG-47), a regulator of cell-autonomous immunity, does not localize to mycobacterial or listerial phagosomes in IFN-gamma-induced mouse cells. J Immunol 191:1765–74.

44. Feng CG, Zheng L, Jankovic D, Bafica A, Cannons JL, Watford WT, Chaussabel D, Hieny S, Caspar P, Schwartzberg PL, Lenardo MJ, Sher A. 2008. The immunity-related GTPase Irgm1 promotes the expansion of activated CD4+ T cell populations by preventing interferon-gamma-induced cell death. Nat Immunol 9:1279–87.

45. Coers J, Gondek DC, Olive AJ, Rohlfing A, Taylor GA, Starnbach MN. 2011. Compensatory T cell responses in IRG-deficient mice prevent sustained Chlamydia trachomatis infections. PLoS Pathog 7:e1001346.

46. Pires D, Marques J, Pombo JP, Carmo N, Bettencourt P, Neyrolles O, Lugo-Villarino G, Anes E. 2016. Role of Cathepsins in Mycobacterium tuberculosis Survival in Human Macrophages. Sci Rep 6:32247.

